# *De Novo* Protein Fold Design Through Sequence-Independent Fragment Assembly Simulations

**DOI:** 10.1101/2022.05.16.492148

**Authors:** Robin Pearce, Xiaoqiang Huang, Gilbert S. Omenn, Yang Zhang

## Abstract

*De novo* protein design generally consists of two steps, including structure and sequence design. However, many protein design studies have focused on sequence design with scaffolds adapted from native structures in the PDB, which renders novel areas of protein structure and function space unexplored. Here we developed FoldDesign to create novel protein folds from specific secondary structure (SS) assignments through sequence-independent replica-exchange Monte Carlo (REMC) simulations. The method was tested on 354 non-redundant topologies, where FoldDesign consistently created stable structural folds, while recapitulating on average 87.7% of the SS elements. Meanwhile, the FoldDesign scaffolds had well-formed structures with buried residues and solvent exposed areas that closely matched their native counterparts. Despite the high fidelity to the input SS restraints and local structural characteristics of native proteins, a large portion of the designed scaffolds possessed global folds that were completely different from natural proteins in the PDB, highlighting the ability of FoldDesign to explore novel areas of protein fold space. Detailed data analyses demonstrated that the major contributions to the successful fold design lay in the optimal energy force field, which contains a balanced set of fragment and secondary structure packing terms, and the REMC simulations, which utilize multiple auxiliary movements to efficiently search the conformational space. These results demonstrate FoldDesign’s strong potential to explore both structural and functional space through computational design simulations that natural proteins have not reached through evolution.

**Significance:** Natural proteins were generated following billions of years of evolution and therefore possess limited structural folds and biological functions. There is considerable interest in *de novo* protein design to generate artificial proteins with novel structures and functions beyond those created by nature. However, the success rate of computational *de novo* protein design remains low, where extensive user-intervention and large-scale experimental optimization are typically required to achieve successful designs. To address this issue, we developed a new automated open-source program, FoldDesign, for *de novo* protein fold design which shows improved performance in creating high fidelity stable folds compared to other state-of-the-art methods. The success of FoldDesign should enable the creation of desired protein structures with promising clinical and industrial potential.

## Introduction

Proteins are important biological molecules that perform the majority of cellular functions in living organisms. Their unique and varied functions are made possible by the diverse structural folds adopted by different protein molecules. However, despite the enormous conformational space available, only a tiny portion appears in nature following billions of years of evolution (1). For example, there have been just under 1,500 protein folds classified in the SCOPe database (2) and studies have indicated that the current PDB is nearly complete, representing the vast majority of natural folds (3, 4). Given the vital importance of proteins to living organisms, there has been growing interest in designing artificial proteins with enhanced functionality beyond the native counterparts. However, many attempts have focused on generating new protein sequences starting from the structures of experimentally solved proteins (5–8). While this may be effective in certain cases, protein design starting from solved structures is severely limited as nature has essentially sampled from an insignificant portion of the structure and function space.

*De novo* protein design, which aims to create not only artificial protein sequences but also novel structures tailored to specific design applications, e.g., with specific fold types or binding pockets, has gained considerable traction in recent years, where approaches such as Rosetta have been applied to design proteins with promising therapeutic potential (9–11), novel ligand-binding activity (12, 13), and complex logical interactions (14). Despite the successes, the approach remains somewhat of an art form, where large-scale experimental optimization is typically required to generate successful designs (9, 11). In particular, extensive user-intervention during scaffold creation and selection is often required during the design simulations (12, 15). Nevertheless, automated fold design tailored to specific applications is highly nontrivial because traditional homologous structure assembly programs often create folds that are similar to the template structures even when distracted with strong external spatial restraints (16, 17). Although *ab initio* structure assembly approaches, such as QUARK (18) and Rosetta (19), can create template-free models, they need to start from specific natural sequences and often create conformations that either converge to specific folding clusters or are not protein-like (20). Anishechenko et al. recently performed an interesting study that combined deep neural-network training with structural refinement simulations to ‘hallucinate’ proteins; the method could create novel protein sequences but the structural folds were generally close to PDB structures (with an average TM-score=0.78) (21). Meanwhile, the resulting protein folds were largely randomized depending on the stochastic process of the design iterations. Thus, development of automated algorithms capable of precisely designing any required fold type with limited human intervention is critical to improve the efficiency and success rate of *de novo* protein design.

Toward this goal, we developed a new automated pipeline, FoldDesign, to create desired protein folds starting from user-specified restraints, such as the secondary structure (SS) topology and/or inter-residue contact and distance maps, through sequence-independent replica-exchange Monte Carlo (REMC) simulations. Since the designed folds do not necessarily have experimental counterparts, we designed several objective assessment criteria based on the satisfaction rate of the input requirements and the folding stability of the designs (see Fig. S1 in the Supporting Information (SI)). The results showed that FoldDesign is capable of producing protein-like structural folds that closely recapitulate the input features with enhanced folding stability, significantly outperforming other start-of-the-art approaches on the large-scale benchmark tests. The online server and standalone package for FoldDesign are freely available to the community at https://zhanggroup.org/FoldDesign/ and https://github.com/robpearc/FoldDesign, respectively.

## Results and Discussion

FoldDesign is an automated algorithm for sequence-independent, *de novo* protein fold design, where the flowchart is outlined in Fig. 1. The program takes as input the SS topology for a designed structure scaffold, which includes the length, order, and composition of the SS elements. A set of structural fragments with lengths between 1-20 residues is then collected from the PDB library by scoring the similarity between the input SS and the SS of the PDB fragments. These fragments are finally assembled together by REMC folding simulations to generate protein-like structural scaffolds that satisfy the input constraints, where the lowest free-energy structure selected by conformational clustering (22) is subjected to further atomic-level refinement to produce the final structural design (see Methods).

**Figure 1.**
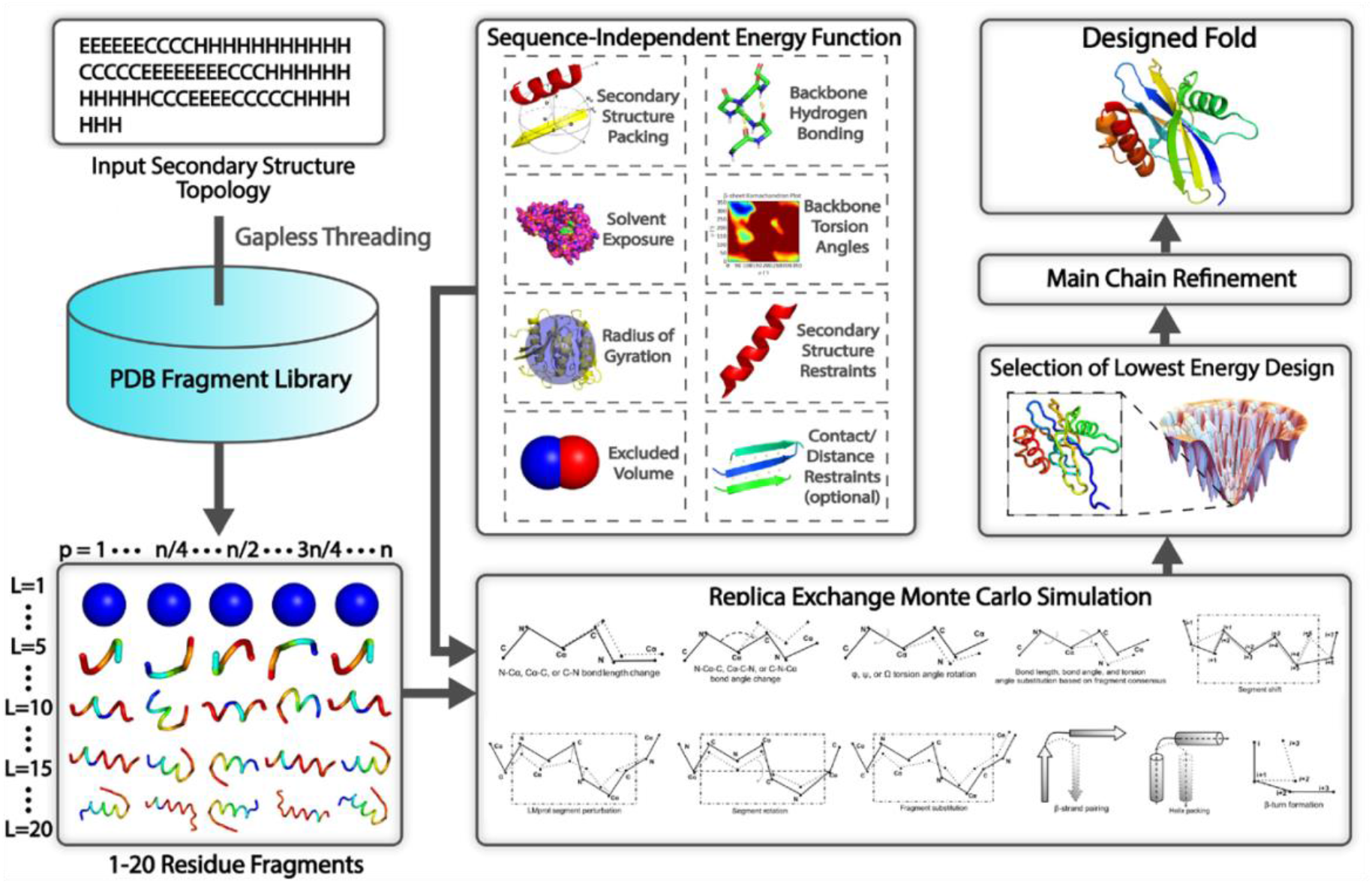
Overview of the FoldDesign pipeline. Starting from a user-defined secondary structure topology as well as any further design constraints such as inter-residue contacts or distances, FoldDesign identifies 1-20 residue structural fragments from the PDB with secondary structures that match the input constraints. These fragments are then assembled together along with 10 other conformational movements during the replica-exchange Monte Carlo folding simulations under the guidance of a sequence-independent energy function that accounts for the fundamental forces that underlie protein folding. The lowest energy structure produced during the folding simulations is selected for further atomic-level refinement by ModRefiner to produce the final designed structure.

### Auxiliary movements improve the folding simulation efficiency and ability to identify low energy states

Fragment substitution is the predominant movement used by FoldDesign, which involves the replacement of a selected region of a decoy structure with the structure from one of the identified fragments collected from the PDB. However, fragment substitution may cause large conformational changes that prevent the movement from being accepted. To improve the simulation efficiency, FoldDesign introduces 10 additional auxiliary movements, including bond length and angle perturbations, segment rotations, torsion angle substitutions, and those that form packing interactions between specific secondary structure elements (see Supplementary Text S1 and Fig. S4).

Fig. 2A displays the FoldDesign energies of the best structures produced for each of the 354 test SS topologies (see Methods), either using the full set of 11 conformational movements or only using fragment substitution. It can be observed that the auxiliary movements enable the simulations to find structures with significantly lower energies than those found using fragment substitution alone. Overall, the average FoldDesign energy of the best structures produced using the full movement set was −529.5 *k_B_T* compared to −449.7 *k_B_T* when using only fragment substitution, where the difference was highly statistically significant with a p-value of 2.1E-66 as determined by a paired two-sided Student’s t-test. In addition to the improved ability to sample low energy states, the auxiliary movements reduced the simulation times required to fold the proteins. Fig. 2B plots the simulation time versus the protein length for each of the test topologies. From the figure, a clear reduction in the simulation time required can be seen across all protein lengths, where the average time for the simulations with the full movement set was 9.6 hours compared to 22.8 hours for the simulations that used only fragment substitution. This reduction in simulation time is due to the fact that fragment substitution is computationally expensive and requires additional loop closure to ensure that it does not cause large downstream perturbations, while the auxiliary movements are comparatively fast.

**Figure 2.**
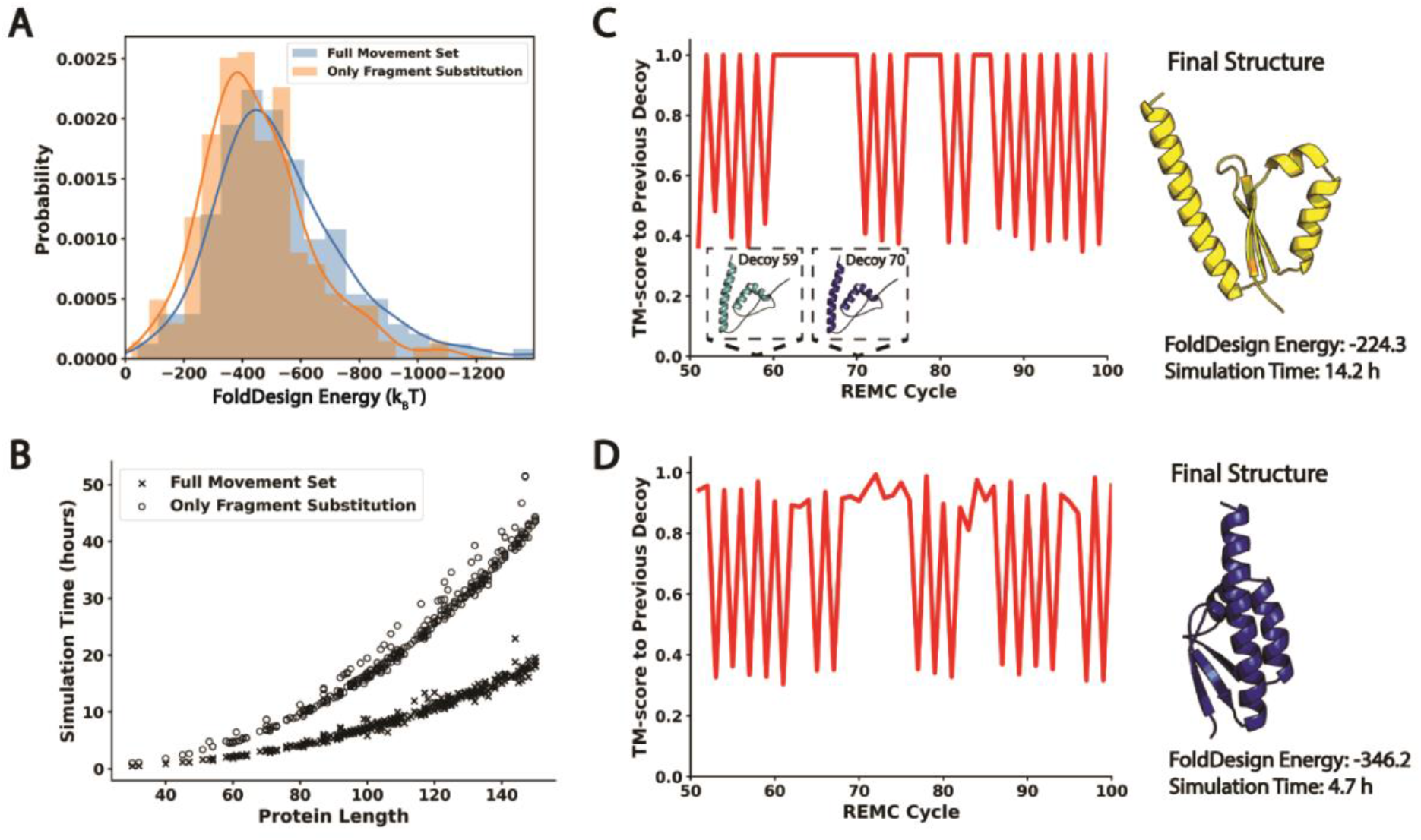
Importance of the auxiliary conformational movements. A) Energy distributions for the designs produced by the FoldDesign simulations using the full movement set and using only fragment assembly. B) Simulation time required versus protein length for FoldDesign using the full movement set and fragment assembly alone. C-D) Two representative case studies that demonstrate the dynamics of the folding simulations without (C) and with (D) the auxiliary movements. The y-axis displays the TM-score between the decoy at REMC cycle *i* compared to the decoy at cycle *i-1*.

In Figs. 2C-D, we further present a representative case study for the topology from the PDB protein 1ec6A, which adopts an α/β fold. Fig. 2C shows the conformational dynamics of the decoys produced during the lowest-temperature replica of the simulations using only the fragment substitution movement, while Fig. 2D uses the full movement set, by plotting the TM-score between the decoy at REMC cycle *i* compared to cycle *i-1* from cycles 50-100. In Fig. 2C, there are several plateaus where no movement could be accepted, leading to identical conformations between a number of the cycles, where the most notable plateau lasted for 11 cycles (cycles 59 through 70). On the other hand, with the full movement set in Fig. 2D, no such plateaus were observed. Although several cycles had very similar structures, which may be caused by subtle conformational refinements such as bond length perturbation, none of the cycles had identical structures. As a result, the simulations using the full movement set generated a structure with an energy of −346.2 *k_B_T* in 4.7 hours compared to a structure with an energy of −224.3 *k_B_T* in 14.2 hours using only fragment substitution.

### FoldDesign scaffolds closely match the input constraints

To assess the ability to perform structural fold design, we list in Table 1 a summary of the FoldDesign results in terms of the average Q3 scores on the 354 test topologies. As a comparison, we list the results from the state-of-the-art Rosetta method (23), which also starts from the desired secondary structure of a designed scaffold, where a detailed description of the procedures used to run Rosetta is given in Supplementary Text S3. Here, the Q3 score is defined as the fraction of positions with secondary structures that are identical to that of the input topology. Following fold generation, the secondary structures of the designed scaffolds for both FoldDesign and Rosetta were assigned using DSSP (24) and compared to the input for each protein.

**Table 1.** Comparison of the Q3 scores for the structures produced by FoldDesign and Rosetta on the 354 test secondary structure topologies. Here, the Q3 score is defined as the fraction of positions in the designed structures whose secondary structures were identical to the input secondary structures. The results are further separated based on the fold type (α, β, and α/β) and the *p*-values were calculated using paired, two-sided Student’s t-tests.

| Method | Q3 Score All<br>($p$-value) | Q3 Score α-proteins<br>($p$-value) | Q3 Score β-proteins<br>($p$-value) | Q3 Score $\alpha/\beta$-proteins<br>($p$-value) |
| --- | --- | --- | --- | --- |
| FoldDesign | **0.877 (\*)** | **0.934 (\*)** | **0.863 (\*)** | **0.875 (\*)** |
| Rosetta | 0.833 (1.7E-08) | 0.828 (5.4E-05) | 0.829 (0.10) | 0.835 (4.5E-06) |

Overall, FoldDesign achieved an average Q3 score of 0.877 compared to 0.833 for Rosetta with a p-value of 1.7E-08. When considering the Q3 scores for α-proteins, β-proteins, and α/β-proteins separately, FoldDesign achieved Q3 scores of 0.934, 0.863, and 0.875, compared to 0.828, 0.829, and 0.835, respectively, for Rosetta. Thus, across all fold types, FoldDesign was able to generate structures that more closely matched the input topologies than Rosetta. This partially reflects the advanced dynamics of the folding simulations as well as the effectiveness of the optimized energy function in FoldDesign.

Although no user-defined distance restraints were included in the above tests, these are still important in many design cases where recapitulation of specific folds is desired. In Table S1, we extracted the pairwise Cα distances from the native structures in the test set and used them as restraints during the design simulations. From the table, it can be seen that FoldDesign was able to generate designs that closely matched the native structures with average TM-scores/RMSDs of 0.993/0.31Å, 0.993/0.27Å, 0.992/0.32Å, and 0.994/0.31Å for all, α, β, and α/β topologies, respectively. Here, TM-score (25) is a structure comparison metric that takes a value in the range (0, 1], where a value of 1 indicates an identical match between two structures and a value ≥0.5 signifies that two proteins share the same global fold (26). Therefore, the FoldDesign structures nearly perfectly recapitulated the desired folds when guided by user-defined distance restraints. Additionally, the mean absolute errors (MAEs) between the Cα distance maps extracted from the designed folds and native structures were 0.148, 0.115, 0.130, and 0.154 Å for all, α, β, and α/β topologies, respectively, confirming that the generated structures closely satisfied the given distance restraints.

### FoldDesign generates low energy, native-like protein structures

While an important metric, the Q3 score is unable to provide a complete picture of the physical quality of the designs. In theory, a method could produce trivial or even unfavorable folds that satisfy the desired secondary structures. Thus, a more detailed analysis of the energetics and physical characteristics of the produced structures had to be performed (Fig. S1). As the designed scaffolds for FoldDesign and Rosetta are both sequence-independent and many of the traditional scoring and assessment tools are sequence-specific, the sequence for each scaffold had to be designed before further quantitative analysis could be conducted. To design the sequences for each scaffold, two sequence design methods were used, EvoEF2 (27) and RosettaFixBB (28), where the backbone structures of the designed scaffolds were kept fixed during the sequence design to ensure a fair comparison of the scaffolds that were directly output by FoldDesign and Rosetta. Here, RosettaFixBB and EvoEF2 are sequence design methods that perform Monte Carlo sampling in sequence space guided by combined physics- and knowledge-based energy functions. 100 sequences were designed for each scaffold, where the average results from the 10 lowest energy sequences were reported for both FoldDesign and Rosetta in the following analyses.

First, Fig. 3A shows that the percent of buried residues for the FoldDesign scaffolds closely resembled the native protein structures from which the input secondary structures were extracted. For example, in the native structures, 19.2% of the residues were buried in the hydrophobic core, compared to 20.2% and 17.2% for the FoldDesign scaffolds whose sequences were designed by EvoEF2 and RosettaFixBB, respectively. However, for Rosetta, only 9.8% and 7.5% of the residues were buried in the hydrophobic core. Additionally, the solvent accessible surface area (SASA) for the native proteins was 7081.8 Å^2^ compared to 6964.9 Å^2^ and 7376.3 Å^2^ for the FoldDesign scaffolds whose sequences were designed by EvoEF2 and RosettaFixBB, while the average SASA for the corresponding Rosetta scaffolds was 8721.2 Å^2^ and 8944.2 Å^2^, respectively. These results indicate that the FoldDesign scaffolds possessed more compact hydrophobic cores and less solvent exposed area than the Rosetta scaffolds and shared a higher similarity to the native structures for these characteristics. The difference is in part due to the fact that FoldDesign includes a number of energy terms that promote the formation of well-packed structures; these include specific fragment-derived distance and solvation potentials, generic backbone atom distance energy terms, and detailed secondary structure packing terms (see Supplementary Text S2). In addition, the energy weights were carefully optimized using the results of the design simulations to ensure the formation of well-folded globular proteins (see Methods).

**Figure 3.**
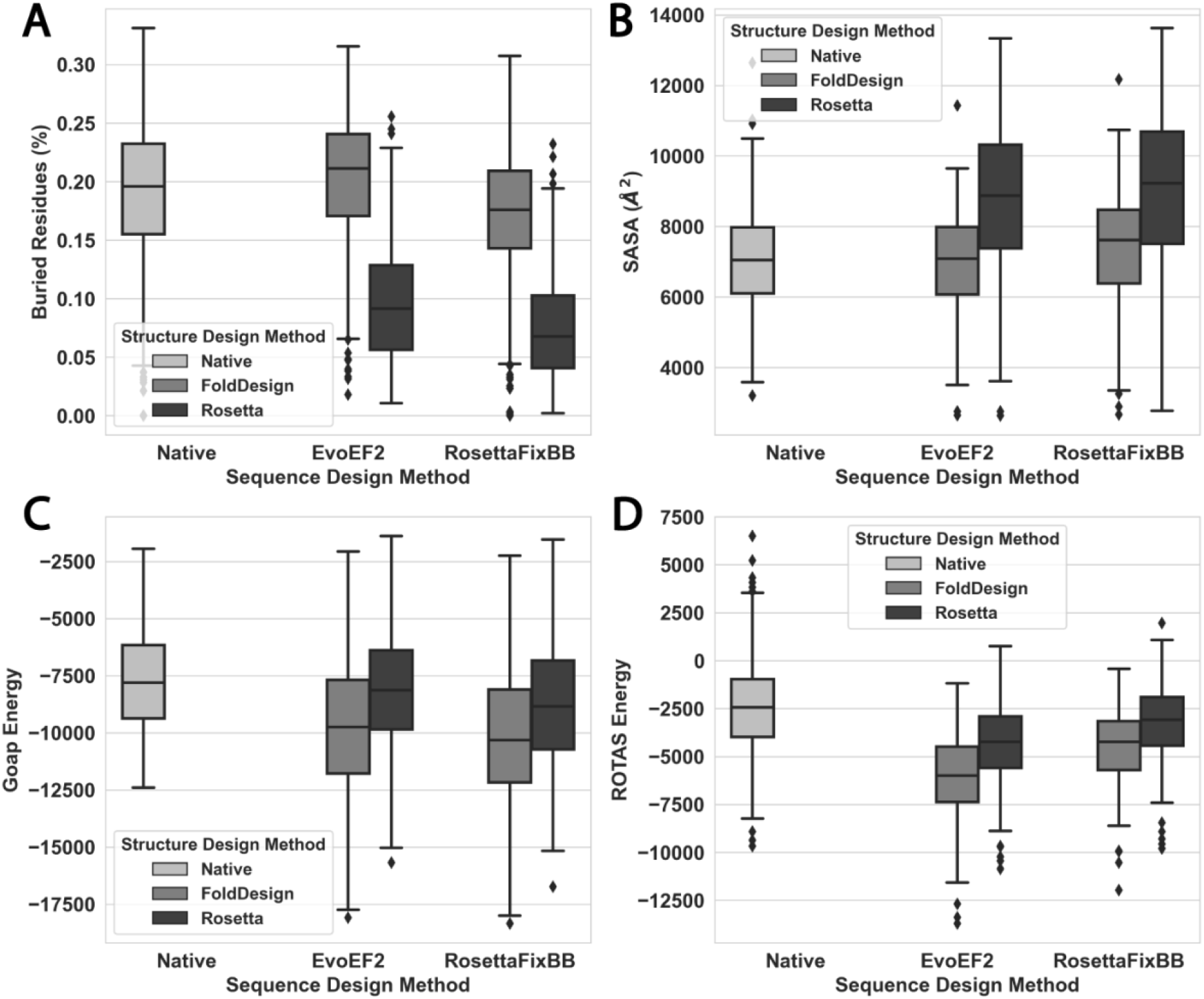
Comparison of the physical characteristics and energies for the designed folds by FoldDesign and Rosetta on the 354 test proteins, where the sequence for each scaffold was designed by EvoEF2 and RosettaFixBB, respectively. The native designation represents the proteins from which the secondary structures of the designed folds were derived. A) Proportion of buried residues is plotted for each protein, where a buried residue was defined as having a relevant solvent accessible surface area <5%. B) Solvent accessible surface area (SASA) for each protein in the test set and those generated by FoldDesign and Rosetta. C-D) Energies for each protein calculated by GOAP and ROTAS.

In Figs. 3C and 3D, we further display the energies of the designed scaffolds by FoldDesign and Rosetta, as assessed by the two leading atomic-level statistical energy functions, GOAP (29) and ROTAS (30). For the sequences designed by EvoEF2 and RosettaFixBB, the FoldDesign scaffolds had average GOAP energies of −9736.9 and −10166.7, respectively, which were significantly lower than the GOAP energies of −8174.5 and −8838.8 for the Rosetta scaffolds, with p-values of 3.4E-13 and 4.3E-10, respectively. Similar trends were observed for ROTAS. For the sequences designed by EvoEF2 and RosettaFixBB, the FoldDesign scaffolds had average ROTAS energies of −6110.3 and −4446.5 compared to −4360.8 and −3281.5 for the corresponding Rosetta designs; the differences were statistically significant with p-values of 6.8E-27 and 1.3E-15. Overall, the FoldDesign scaffolds possessed more tightly packed hydrophobic cores and were energetically more favorable than the Rosetta scaffolds, with GOAP energies that were 19.1% and 15.0% lower than the Rosetta scaffolds and ROTAS energies that were 40.1% and 35.5% lower than the Rosetta scaffolds depending on the sequence design method that was used, although both folding simulations did not use any of the third-party energy functions for optimization.

### FoldDesign generates stable structures with novel folds

To further assess the stability of the designed structures, molecular dynamic (MD) simulations were run starting from the designed scaffolds by FoldDesign and Rosetta. MD is a useful tool as it allows for the study of protein motion and stability beyond static measurements such as energy calculations, where 20 ns unconstrained MD simulations were carried out using GROMACS (31) with the CHARMM36 force field (see Methods). To determine the stability of the structures, the TM-scores (25) between the initial designed scaffolds and the final MD structures were calculated, where the results are depicted in Figs. 4A-B for the structures whose sequences were designed by EvoEF2 and RosettaFixBB, respectively.

**Figure 4.**
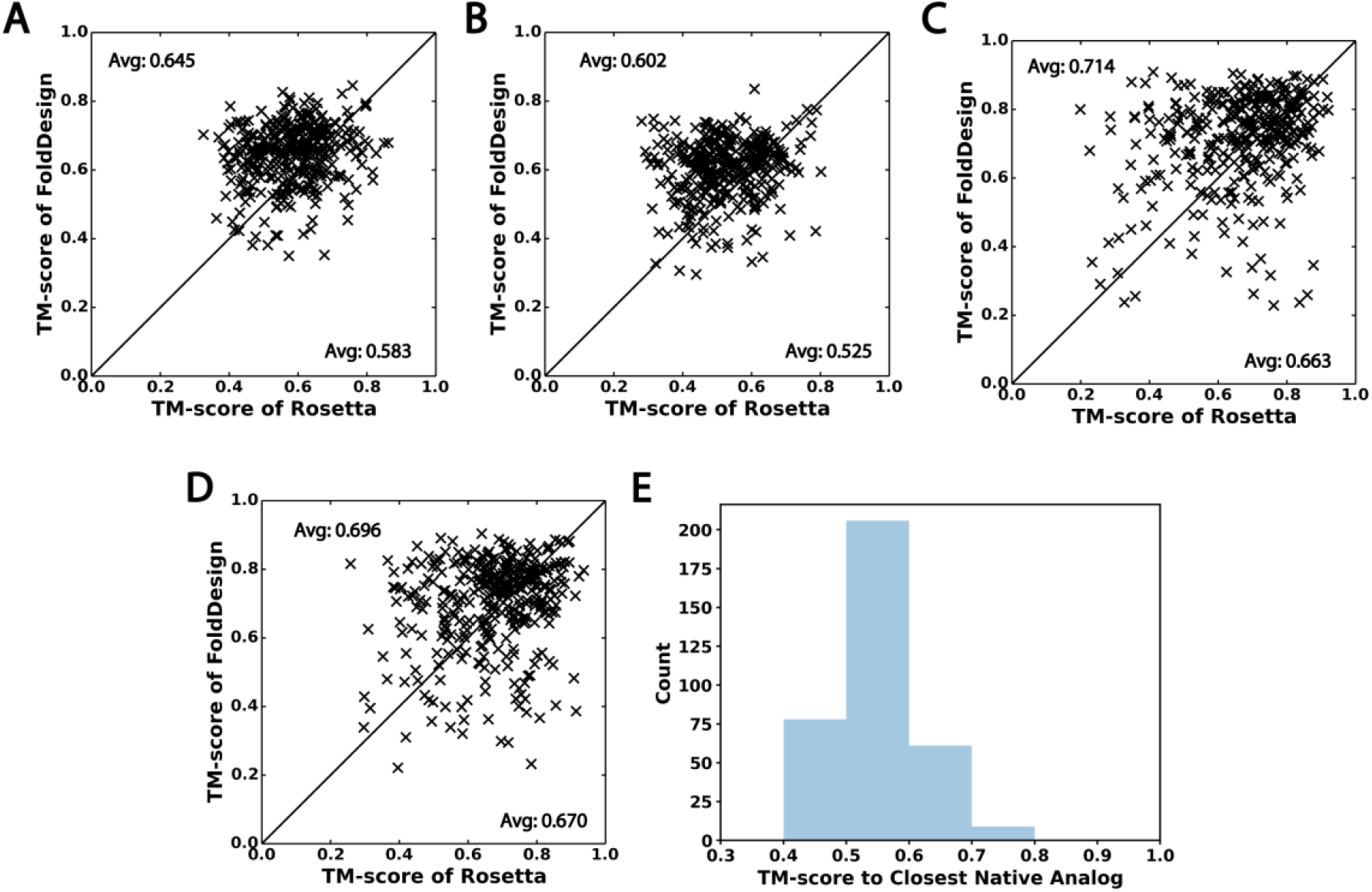
Analysis of the FoldDesign and Rosetta scaffolds using molecular dynamics (A-B) and protein structure prediction by AlphaFold2 (C-D). A-B) TM-scores of the FoldDesign and Rosetta scaffolds relative to their final structures following 20 ns MD folding simulations, where the sequence for each scaffold was designed by EvoEF2 (A) and RosettaFixBB (B). C-D) TM-scores of the FoldDesign and Rosetta scaffolds relative to the structures predicted by AlphaFold2 starting from the EvoEF2 (C) and RosettaFixBB (D) sequences designed for each scaffold. E) TM-score distribution between the FoldDesign structures and their closest native analogs obtained by searching the designed scaffolds through the PDB using TM-align.

From the figures, it can be seen that the TM-scores between the initial FoldDesign scaffolds and the final MD structures were higher than those for the Rosetta scaffolds, indicating a closer match and thus more stable conformations for the FoldDesign scaffolds. For instance, the average TM-score between the FoldDesign scaffolds and final MD structures for the EvoEF2 sequence designs was 0.645 compared to 0.584 for the corresponding Rosetta scaffolds (Fig. 4A), where the difference was statistically significant with a p-value of 7.4E-19. A similar trend was observed for the scaffolds whose sequences were designed by RosettaFixBB, where the average TM-score between the initial FoldDesign structures and the final MD structures was 0.602 compared to 0.525 for the Rosetta scaffolds, corresponding to a p-value of 4.6E-26 (Fig. 4B). Furthermore, when considering a cutoff TM-score of 0.5, 93.7% and 87.9% of the FoldDesign scaffolds whose sequences were designed by EvoEF2 and RosettaFixBB, respectively, shared the same global folds as their final MD structures, compared to 77.1% and 54.8% of the corresponding Rosetta structures. Fig. 5A shows three examples selected from among the most stable FoldDesign scaffolds, where the TM-scores were all greater than 0.8 and the RMSDs were less than 2 Å, indicating a close atomic match between the designed scaffolds and the final MD structures. Overall, the vast majority of the FoldDesign scaffolds possessed stable global folds, outperforming the state-of-the-art Rosetta protocol across the test set.

**Figure 5.**
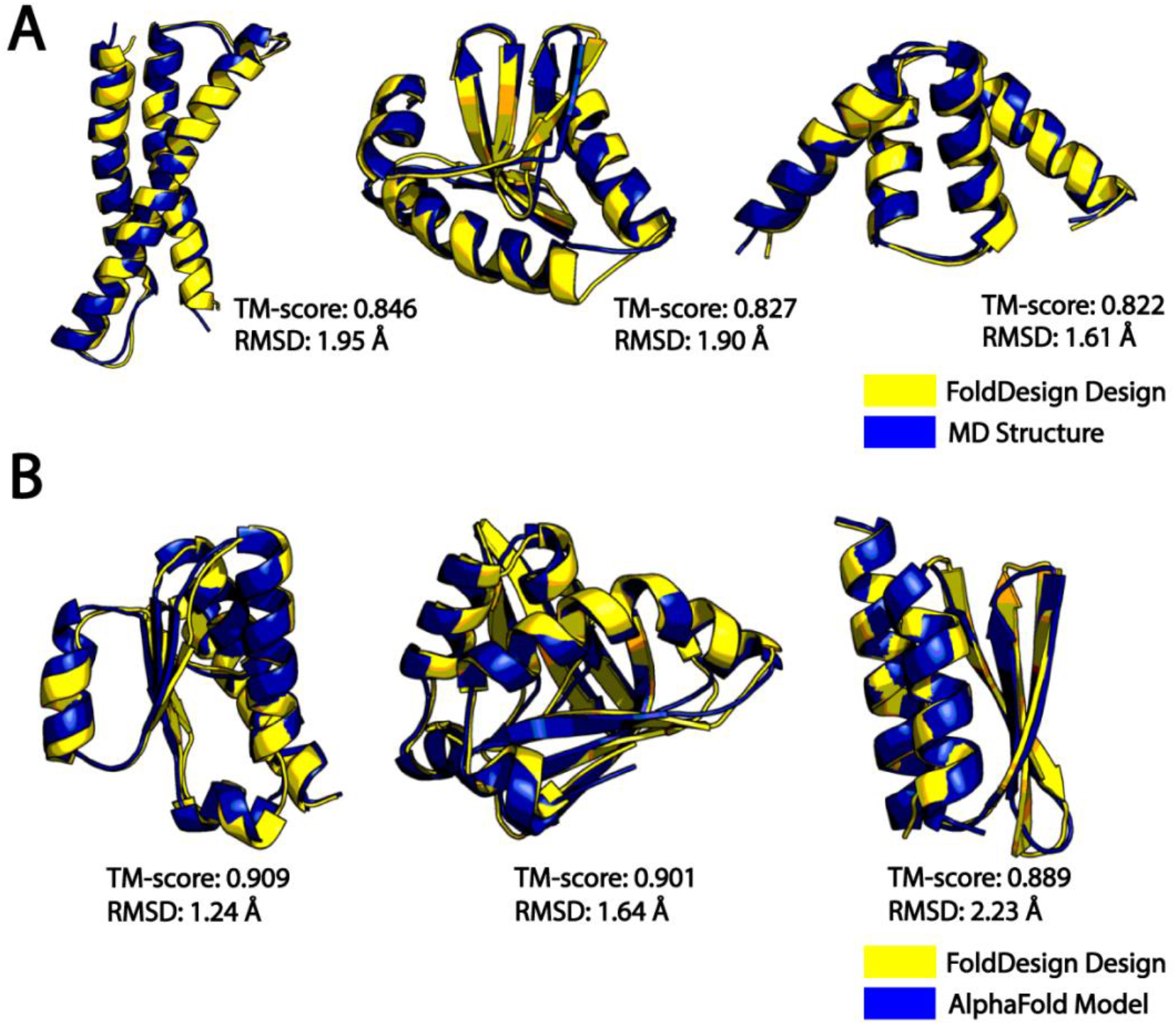
Examples of stable, well-folded FoldDesign scaffolds as assessed by molecular dynamics (A) and AlphaFold2 (B), where the sequences for each scaffold were designed by EvoEF2. A) The initial FoldDesign structures (yellow) superposed with the final MD structures (blue). B) The FoldDesign scaffolds (yellow) superposed with the AlphaFold2 models (blue).

Interestingly, despite the high fold stability with local structural features that were highly similar to the native proteins, a large portion of the FoldDesign scaffolds adopted novel folds that were different from what exists in the PDB. Fig. 4E presents the histogram distribution of TM-scores between the FoldDesign scaffolds and the closest structures identified by TM-align (32) from the PDB, where the average TM-score of 0.551 was relatively low given the searching power of TM-align and the near completeness of the PDB (3, 32). Of the 354 designs, 79 had a TM-score below 0.5 to any structure in the PDB, indicating they possessed novel folds. Furthermore, 74 of the 79 novel structures whose sequences were designed by EvoEF2 had stable global folds with TM-scores >0.5 to their final structures output by the MD simulations. Moreover, there was no obvious difference between the novel folds and other folds in terms of stability, as the TM-score distributions between the designs and the final MD structures were quite similar for them (Fig. S3), where their average TM-scores were 0.647 and 0.645, respectively. These results demonstrate that FoldDesign is capable of producing stable scaffolds, while allowing for the exploration of novel areas of protein fold space.

### Protein structure prediction indicates FoldDesign produces well-folded structures

As additional proof of the foldability of the designed structures, we examined the structural similarity between the designed scaffolds and the predicted models generated by the state-of-the-art AlphaFold2 structure prediction program (33) starting from the designed sequences for each scaffold. As protein structure prediction is essentially the inverse problem of protein design, it would stand to reason that well-formed structure designs should be able to be recapitulated starting from their corresponding designed sequences. Given that AlphaFold2 is a deep learning-based modeling program, its performance largely depends on collecting meaningful MSAs (33). Thus, since *de novo* designed proteins almost always lack natural sequence homologs, to remove the bias in MSA collection, we constructed the input MSAs by taking the 100 sequences designed by EvoEF2 and RosetaFixBB for each of the FoldDesign/Rosetta scaffolds.

As shown in Table S2, when starting from the sequences designed by EvoEF2, the average TM-score between the AlphaFold2 models and the FoldDesign scaffolds was 0.714 compared to 0.663 for the Rosetta scaffolds, where the difference was statistically significant with a p-value of 4.6E-09. In Figure 3C, we present a head-to-head TM-score comparison, where the FoldDesign scaffolds had higher TM-scores than the corresponding Rosetta scaffolds for 211 cases, while Roseta did so for 133 of the 354 cases. If we consider the number of cases with TM-score ≥0.5, 324 (or 91.5%) of the FoldDesign scaffolds shared the same global folds as the AlphaFold2 models compared to 315 (or 90.0%) of the scaffolds by Rosetta. These results demonstrate that the FoldDesign scaffolds more closely resembled the AlphaFold2 models than the Rosetta scaffolds did, indicating their enhanced stability/foldability. Similar patterns were observed for the sequences designed by RosettaFixBB, where the average TM-score between the FoldDesign scaffolds and AlphaFold2 models was 0.696 compared to 0.670 for Rosetta with a p-value of 3.0E-04 (Table S2). Moreover, 208 of the 354 FoldDesign scaffolds had higher TM-scores than the Rosetta scaffolds and 315 (or 89.0%) of the designs had TM-scores ≥0.5 (Fig. 4D).

Fig. 5B presents three examples from some of the closest matches between the FoldDesign scaffolds and AlphaFold2 models, where each had a TM-score greater than or close to 0.9 and RMSDs below 2.25 Å, indicating a close atomic match between the designed scaffold and the predicted models. From these results, it can be seen that the FoldDesign scaffolds more closely matched the predicted models than the Rosetta scaffolds did, and the overwhelming majority of the designs shared the same global folds as the AlphaFold2 models. This structural consistency may suggest that FoldDesign captures some structural characteristics that have been integrated in the AlphaFold2 learning process.

## Concluding Remarks

Protein design generally consists of two steps, including structural fold design and sequence design. Many protein design efforts have focused on the second step of sequence design with input scaffolds taken from existing protein structures in the PDB. Despite the success, such experiments constrain design cases to the limited number of folds adopted by natural proteins, while curtailing the exploration of novel areas of protein structure and biological function.

In this work, we developed a pipeline, FoldDesign, for *de novo* protein fold design. Unlike traditional protein folding simulations which start from native sequences and therefore, as expected, often result in folds that are similar to what exists in the PDB library, FoldDesign starts from structural restraints (e.g., secondary structure assignments and/or inter-residue distance restraints) and performs folding simulations under the guidance of an optimized sequence-independent energy function. Large-scale tests on a set of 354 unique fold topologies demonstrated that FoldDesign could create protein-like folds with closer Q3 score similarity to the desired structural restraints than the state-of-the-art Rosetta design program. Meanwhile, the FoldDesign scaffolds had well-compacted core structures with buried residues and solvent exposed areas that closely resembled natural proteins, while MD simulations showed that the folds were more stable than those produced by Rosetta. Importantly, FoldDesign is capable of designing folds that are completely different from the native structures in the PDB, highlighting its ability to explore novel areas of protein structure space despite the high fidelity to the input restraints and the native local structural characteristics. Detailed data analyses showed that one of the major contributions to the success of FoldDesign lies in the optimal energy force field, which contains a balanced set of energy terms accounting for fragment and secondary structure packing. In addition, the conformational space is effectively explored by REMC simulations assisted by a composite set of efficient movements.

Although the FoldDesign server outputs both the designed fold and the lowest energy sequence produced by the EvoDesign program, the validation of the designed sequences remains to be experimentally examined. Nevertheless, the studies presented have shown that FoldDesign can be used as a robust tool for generating high-quality, stable structural folds when applied to the very challenging task of completely *de novo* scaffold generation without human-expert interventions. This therefore suggests a strong potential for experimental protein design to effectively explore both structural and functional spaces which natural proteins have not reached through billions of years of evolution.

## Methods

FoldDesign aims to automatically design desired protein structure folds starting from user-specified rules such as secondary structure composition and/or inter-residue contact and distance maps. The pipeline consists of three main steps, including fragment generation, REMC folding simulations, and main chain refinement and fold selection (see Fig. 1).

### Fragment generation

Starting from a user-specified secondary structure, high-scoring fragments are identified from a fragment library, which consists of fragments collected from a non-redundant set of 29,156 high-resolution PDB structures. Gapless threading through the library is performed to generate 1-20 residue fragments, where the fragments are scored based on the compatibility of their torsion angles and secondary structure similarity to the desired secondary structure at each position. The top 200 fragments are generated for each overlapping 1-20 residue window. The information for each fragment includes the backbone bond lengths, bond angles, and torsion angles, as well as other useful data such as the position-specific solvent accessibility and Cα coordinates, which are later used to derive distance and solvation restraints.

### REMC folding simulations and refinement

Following fragment generation, REMC folding simulations are performed in order to assemble full-length structural models, where each simulation uses 40 replicas and runs 500 REMC cycles (see Text S1 for a full description of the REMC parameters and movements). The protein conformation in FoldDesign is represented with a course-grained model, which specifies the backbone N, Cα, C, H, and O atoms as well as the Cβ atoms and an atom that represents the side-chain center of mass (Fig. S2). To allow for a less biased exploration of structure space, the energy terms used by FoldDesign are sequence-independent, where the side-chain center of mass for Valine is used as the generic center of mass for each residue to minimize steric clashes.

The initial conformations are produced by randomly assembling different high-scoring 9 residue fragments and then minimized using a set of 11 movements. Here, the major conformational movement is fragment substitution, which involves swapping a selected region of a decoy structure with the structure from one of the fragments randomly selected from the fragment library. Next, cyclical coordinate descent loop closure (34) is used to minimize the structural perturbations downstream. Since FoldDesign uses 1-20 residues fragments, larger fragment insertions are typically attempted during the initial REMC cycles, while smaller ones are attempted during the later steps of the simulations to improve its acceptance rate when the protein is more globular and well-folded. In addition to fragment insertion, 10 other conformational movements are attempted throughout the course of the simulations, including perturbing the backbone bond lengths, angles or torsion angles, segment rotations, segment shifts, and movements that form specific interactions between different secondary structure elements, where these are described in Supplementary Text S1 and Fig. S4.

The movements are accepted or rejected using the Metropolis criterion (35), where the energy for each conformation is assessed by the following energy function:

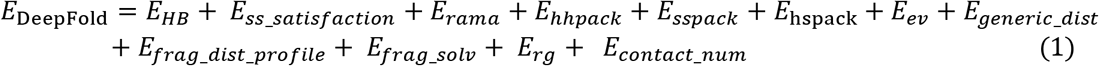

Here, *E_HB_*, *E_ss_satisfaction_*, *E_rama_*, *E_hhpack_*, *E_sspack_*, *E*_hspack_, *E_ev_*, *E_generic_dist_*, *E_frag_dist_profile_*, *E_frag_solv_*, *E_rg_*, and *E_contact_num_* are terms that account for backbone hydrogen bonding, the satisfaction rate of the input secondary structure, Ramachandran torsion angles, helix-helix packing, strand-strand packing, helix-strand packing, excluded volume, generic backbone atom distances, fragment-derived distance restraints, fragment-derived solvent accessibility, radius of gyration, and expected contact number, respectively. A more detailed explanation of these terms is given in Supplementary Text S2. After the REMC simulations are completed, the design with the lowest energy is selected for further atomic-level refinement, for which sequence design and structural refinement are performed iteratively using EvoDesign (5) and ModRefiner (36), respectively.

### Training and test dataset collection

To test FoldDesign’s ability to perform *de novo* protein fold design, we collected a non-redundant set of secondary structure sequences. This was accomplished by extracting the 3-state secondary structures from 76,166 protein domains in the I-TASSER template library (37, 38) using DSSP (24). All of the pairwise secondary structure alignments were obtained using Needleman-Wunsch dynamic programming to align the 3-state secondary structure sequences. The target sequences were then clustered based on the distance matrix defined by their secondary structure identities, i.e., the number of identical secondary structures divided by the total alignment length, where an identity cutoff=70% was used to define the clusters.

The identified clusters were further refined by eliminating atypical secondary structure topologies (clusters with less than 10 members) and by selecting only those clusters where a clear relationship existed between the secondary structure and the tertiary structure adopted by the cluster members. The latter requirement was accomplished by using TM-align (32) to perform structural alignment between each cluster member and the cluster center, where conserved clusters were required to have an average TM-score ≥0.5 between the members and cluster center. Finally, we obtained 461 clusters; 107 and 354 secondary structure sequences were used for the training and test sets, respectively. The training set was composed of 22 α, 25 β, and 60 α/β topologies, while the test set was composed of 24 α, 55 β, and 275 α/β topologies.

### FoldDesign energy function optimization

In order to ensure proper structure generation, each energy term must be carefully weighted in the FoldDesign energy function. This was done on the 107 training topologies. Briefly, a grid searching strategy was used to optimize the weights, where all weights were initially assigned as 0, except for the weight for the steric clash term, which was set to 1.0. Then the values for each weight were adjusted one-at-a-time around the grid values and the FoldDesign simulations were run to produce scaffold structures using the new weight set. After structure generation, the sequences for each scaffold were designed using EvoEF2 (27) and the designed structures were assessed based on:

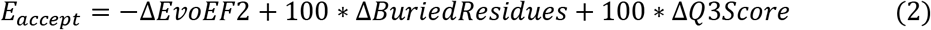

where, Δ*EvoEF2*, Δ*BuriedResidues*, and Δ*Q3Score* are the changes in the average EvoEF2 energy, percent of buried residues, and secondary structure Q3 score, respectively, between the structures produced by the old and new weight sets. If the new weighting parameter increased the value of *E_accept_*, the weights were accepted. Once the initial weights for each energy term were determined, many more iterations were conducted to precisely fine-tune their values based on Eq. (2) as well as by hand inspection of the structures. Although time-consuming, the process of directly optimizing the weights based on the results of the folding simulations resulted in high quality scaffolds with physical characteristics that resembled native proteins.

### Molecular dynamics simulations for examining fold stability

To examine the stability of the FoldDesign scaffolds, we performed MD simulations starting from the designed structures. For each simulation, a dodecahedron box was constructed with a distance of 10 Å from the solute and filled with TIP3P water molecules, where Na^+^ and Cl^-^ ions were used to neutralize the charge of the system. Following this, energy minimization was carried out using steepest descent minimization with a maximum force of 10 kJ/mol. The system was then equilibrated at 300 K using 100 ps NVT simulations and 100 ps NPT simulations with position restraints (1000 kJ/mol) on the heavy atoms of the protein. After the two equilibration phases, the system was well-equilibrated at the desired temperature and pressure, and unconstrained MD simulations were performed at 300 K for 20 ns. During the simulations, non-bonded interactions were truncated at 12 Å and the Particle Mesh Ewald methods was used for long-range electrostatic interactions. Lastly, the velocity-rescaling thermostat and Parrinello-Rahman barostat were used to couple the temperature and pressure, respectively. 1000 structures were collected from the MD trajectories during the final nanosecond of the simulations. This ensemble was then clustered using the GROMOS method with an RMSD cutoff of 2 Å, and the final MD structure for each simulation was collected from the cluster center.

## Acknowledgements

This work used the Extreme Science and Engineering Discovery Environment (XSEDE (39)), which is supported by the National Science Foundation (ACI-1548562). This work was supported in part by the NIGMS (GM136422, S10OD026825), the NIAID (AI134678), and the NSF (IIS1901191, DBI2030790, MTM2025426) and NCI (U24CA210967) and NIEHS (P30ES017885).

## Author Contributions

Y.Z. conceived and designed the research; R.P. developed FoldDesign, performed the experiments, analyzed the data, and developed the stand-alone package; X.H. assisted with the MD simulations; R.P., X.H., G.S.O., and Y.Z. wrote the manuscript.

