## Supplemental Information for "*De Novo* Protein Fold Design Through Sequence-Independent Fragment Assembly Simulations"

#### Table of Contents

**Supporting Tables**

- **Table S1.** Results of FoldDesign starting from distance restraints extracted from the native structures.
- **Table S2.** Comparison of the AlphaFold models starting from the designed sequences with the FoldDesign and Rosetta designed scaffolds.
- **Table S3.** Feature values for each hydrogen bonding restraint type.

**Supporting Figures**

- **Figure S1.** Benchmark criteria used to assess the designed structures.
- **Figure S2.** Depiction of the reduced model used to represent protein conformations during the FoldDesign simulations.
- **Figure S3.** Assessment of the stability of the novel folds generated by FoldDesign.
- **Figure S4.** Depiction of the conformational movements used by FoldDesign.
- **Figure S5.** Illustration of the features used to calculate the energy for packing two secondary structure elements.

**Supporting Texts**

- **Text S1:** Replica-exchange Monte Carlo simulation parameters and movements.
- **Text S2:** FoldDesign energy function.
- **Text S3:** Rosetta protocol used to generate designed folds.

**References**

### Supplementary Tables

**Table S1.** Results of FoldDesign starting from distance restraints extracted from the native structures. All metrics were computed between the designed and native structures. Here, MAE is the mean absolute error between the Cα distance maps from the designed and native structures and is calculated by $MAE = \frac{\sum_{i=1}^{n}|x_i - y_i|}{n}$, where $x_i$ is a distance from a designed structure, $y_i$ is the corresponding distance from the native structure, and $n$ is the number of considered distances.

| Protein Type | MAE (Å) | TM-score | RMSD (Å) |
|---|---|---|---|
| All | 0.148 | 0.993 | 0.31 |
| α | 0.115 | 0.993 | 0.27 |
| β | 0.130 | 0.992 | 0.32 |
| α/β | 0.154 | 0.994 | 0.31 |

**Table S2.** Results of AlphaFold2 modeling starting from the designed sequences for the FoldDesign and Rosetta scaffolds. *P*-values were calculated using paired, two-sided Student's t-tests.

| Method | TM-score (*p*-value) | RMSD (*p*-value) | # TM-score ≥ 0.5 |
|---|---|---|---|
| ***Sequences designed by EvoEF2*** | | | |
| FoldDesign | **0.714 (*)** | **3.66 (*)** | **324** |
| Rosetta | 0.663 (1.1E-07) | 5.10 (4.6E-09) | 301 |
| ***Sequences designed by RosettaFixBB*** | | | |
| FoldDesign | **0.696 (*)** | **4.13 (*)** | **315** |
| Rosetta | 0.670 (0.004) | 4.95 (3.0E-4) | 310 |

**Table S3.** Feature values for each hydrogen bonding restraint type, $T_k$. The features are presented as averages/standard deviations.

| Restraint Type | Secondary Structure | $f_1$: $D(O_i, H_j)$ (Å) | $f_2$: $A(C_i, O_i, H_j)$ (degrees) | $f_3$: $A(C_i, O_i, H_j)$ (degrees) | $f_4$: $T(C_i, O_i, H_j, N_j)$ (degrees) |
|---|---|---|---|---|---|
| $T_1$ | Helix, $j = i + 4$ | 2.00/0.53 | 147/10.58 | 159/11.25 | 160/25.36 |
| $T_2$ | Helix, $j = i + 3$ | 2.85/0.32 | 89/7.70 | 111/8.98 | -160/7.93 |
| $T_3$ | Parallel Strand | 2.00/0.30 | 155/11.77 | 164/11.29 | 180/68.96 |
| $T_4$ | Antiparallel Strand | 2.00/0.26 | 151/12.38 | 163/11.02 | -168/69.17 |

#### Supplementary Figures

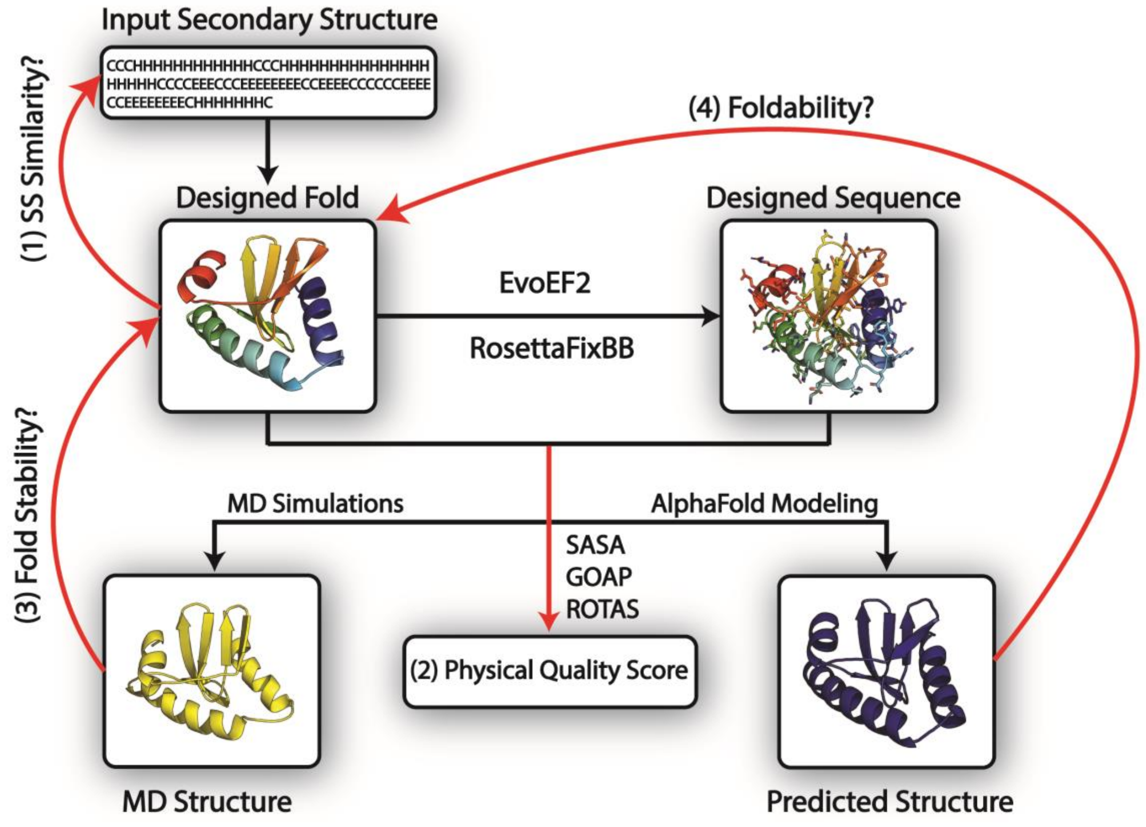


**Figure S1.** Illustrations of the strategies used to evaluate the quality of the FoldDesign scaffolds. The red lines mark the four criteria used to assess the FoldDesign scaffolds: (1) the secondary structure similarity between the input secondary structures and the secondary structures of the scaffolds designed by FoldDesign; (2) the physical quality score including hydrophobic core formation and statistical energies; (3) the fold stability assessed by the structural similarity (TM-score/RMSD) between the FoldDesign scaffolds and the final models after constraint-free molecular dynamic simulations (MD); (4) the foldability as determined by the structural similarity between the FoldDesign scaffolds and the predicted models by AlphaFold2.

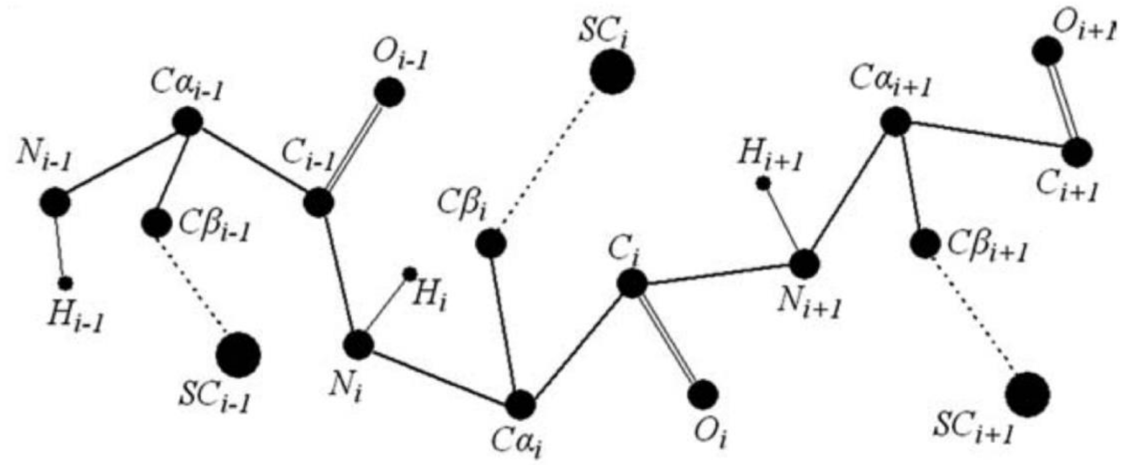


**Figure S2.** Depiction of the reduced model used to represent protein conformations during the FoldDesign simulations, including the backbone atoms (N, H, Cα, C, and O) as well as the Cβ atoms and side-chain centers of mass (SC). The center of mass for Valine is used in this study to evaluate steric clashes.

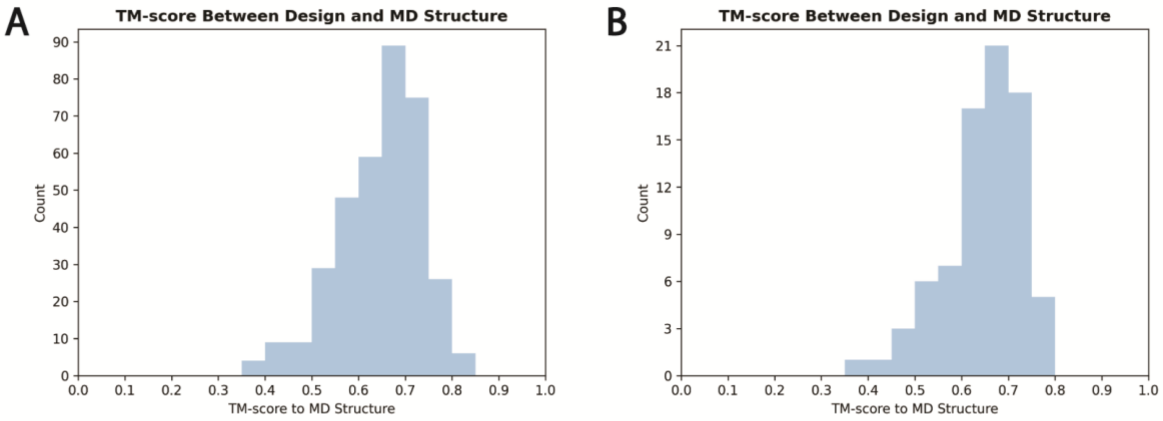


**Figure S3.** Assessment of the stability of the novel folds generated by FoldDesign. A) TM-score distribution between the FoldDesign scaffolds and their final MD structures on the 354 test topologies. B) TM-score distribution between the 79 novel FoldDesign structures and their final MD structures.

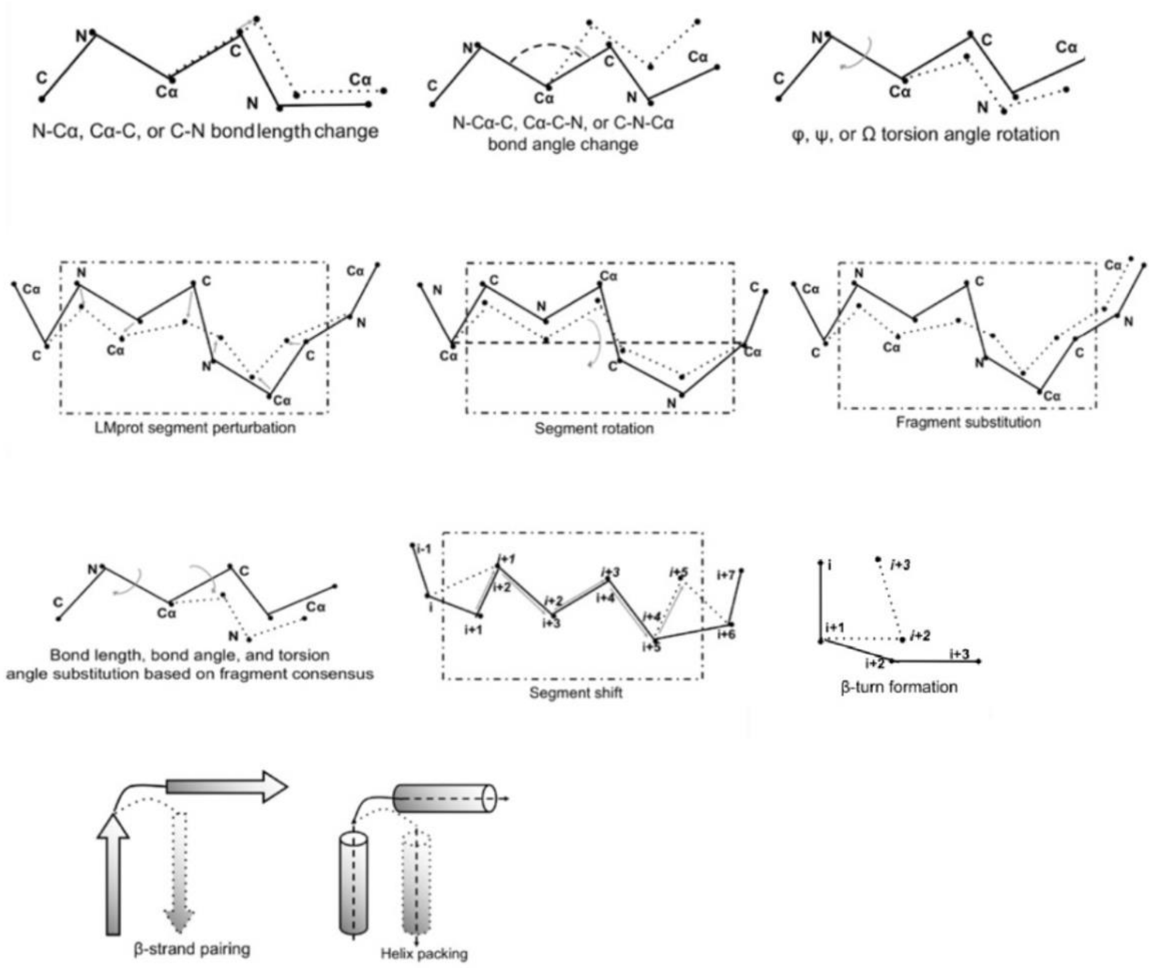


**Figure S4.** Depiction of the conformational movements used by FoldDesign, with explanations in Supplementary Text S1.

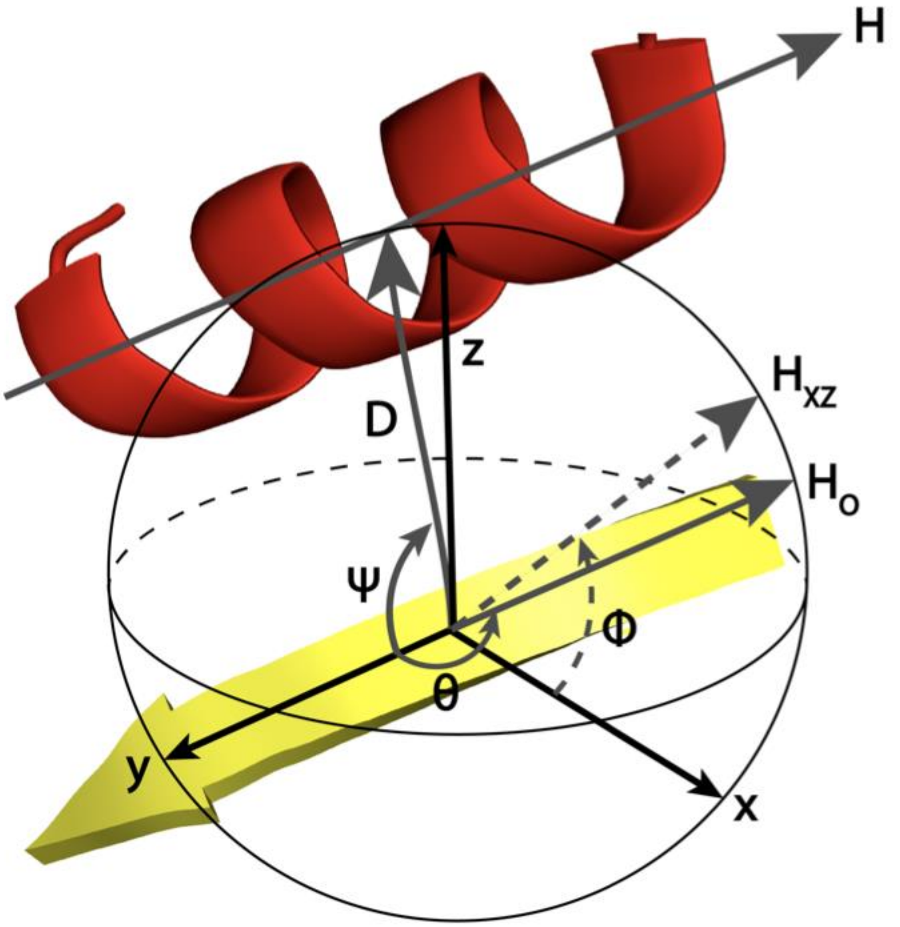


**Figure S5.** Illustration of the features used to calculate the energy for packing two secondary structure elements. Note, here a helix and strand are used, but the parameters are the same for two helices or two strands. The y-axis is defined along the direction of the strand, where the origin is set at the center. D is a vector that represents the distance between the center of the strand and the center of the helix, and the x-axis is defined as the cross product between the y-axis vector and the D vector. The z-axis is defined as the cross product of the y-axis and the x-axis. H is the helical axis and $H_0$ is the helical axis translated to the origin. $H_{xz}$ is the projection of $H_0$ onto the xz-plane. Lastly, $\psi$, $\phi$, and $\theta$ are the angles between the y-axis and the D vector, the x-axis and $H_{xz}$, and the y-axis and $H_0$, respectively.

#### Supplementary Texts

**Text S1: Replica-exchange Monte Carlo simulation parameters and movements.**

The conformational landscape is explored in FoldDesign using replica-exchange Monte Carlo (REMC) simulations. Within REMC, four parameters need to be carefully considered. First, the highest temperature ($T_{max}$) should be high enough to enable the simulation to overcome energy barriers, while the lowest temperature ($T_{min}$) should be low enough to ensure the simulation sufficiently scans the low-energy states. Second, the number of replicas ($N_{rep}$) should be large enough to ensure sufficient chances for the adjacent replicas to communicate with each other. Third, the number of local movements ($N_{sweep}$) before the global swaps should be selected to make the local Metropolis search achieve satisfactory equilibrium. After successive rounds of optimization, the final parameters were selected as: $T_{max} = min(20 * (1 + (L - 100) * 0.004), 20 * 2.5)$, $T_{min} = max(1 * (1 + (L - 100) * 0.001), 1 * 0.5)$, $N_{rep}$ = 40, and $N_{sweep}$ = $30*\sqrt{L}$, where L is the sequence length. A total of 500 REMC simulation cycles are carried out for each design.

During the REMC simulations, 11 different conformational movements are used by FoldDesign, as show in Fig. S4, to sample from structural space. Movements are accepted or rejected using the Metropolis Criterion (1) based on the associated changes in energy calculated by the energy function described in Text S2. The major conformational movement is fragment substitution, where the decoy conformation in a selected region of the protein is replaced with the conformation from one of the highest scoring fragments. In order to perform this movement, it is first necessary to identify local fragments from a fragment library that match the input secondary structure topology. The fragment library is composed of 1-20 fragments from 29,156 high-resolution PDB structures. The information present for each fragment include the position-wise backbone torsion angles $(\phi, \psi, \omega)$, secondary structure, bond lengths, bond angles, solvent accessibility and Cα coordinates. During the movement, the backbone torsion angles $(\phi, \psi, \omega)$ and backbone bond lengths and angles in the decoy structure are swapped with those present in the selected fragment. Next, cyclical coordinate descent loop closure (2) is used to connect the anchor points and prevent large downstream perturbations. Larger insertions are attempted at the beginning of the simulation, when the protein is largely unfolded, and smaller insertions are attempted as the protein become more compact.

In addition to fragment assembly, FoldDesign uses 10 auxiliary movements. The first of the auxiliary movements involves changing the length of one of the backbone bonds by a random value in the range [-0.24 Å, 0.24Å], including the N-Cα, Cα-C, or C-N bonds. The second movement involves randomly changing one of the backbone angles by a value in the range [-10°, 10°], including the $N_i$-$C\alpha_i$-$C_i$, $C\alpha_i$-$C_i$-$N_{i+1}$, and $C_i$-$N_{i+1}$-$C\alpha_{i+1}$ angles, where *i* corresponds to the residue position. The third auxiliary movement changes one or more of the backbone torsion angles $(\phi, \psi, \omega)$. The $\phi$ and $\psi$ angles are updated by sampling from the allowed regions in the Ramachandran plots based on the input secondary structure at a given position. The $\omega$ angle is changed by a value randomly selected from the range [-8°, 8°], where the movement is automatically rejected if it would result in the $\omega$ angle falling outside of the range of (170°, 190°). The fourth movement is LMProt perturbation (3), which randomly changes the positions of the backbone atoms in a selected region and then attempts to restrict all bond lengths and bond angles to physically allowable values. The fifth movement is segment rotation, which rotates the backbone atoms by a random value in the range of (-90°, 90°) for a 2-12 residue segment along the axis defined by the Cα atoms of the first and last residues of the selected region. The sixth movement is similar to the fragment substitution movement but is based on fragment consensus from the 10 residue long fragments. To perform this movement, the 10 residue long identified fragments are clustered based on the distance matrix defined by their $\phi/\psi$ angle pairs. Then during the simulations, the $\phi/\psi$ angle pairs for a 10 residue segment in the decoy structure are swapped for the corresponding angles from the consensus fragments. The seventh movement is a segment shift. It involves shifting the residue numbers in a segment forward or backwards by one residue, which means that the coordinates of each residue are copied from their preceding or subsequent residues in the segment. We then delete the unused coordinates of one residue at the selected terminal region and insert new coordinates for another residue at the other terminal based on physically allowable bond lengths and angles. This movement can easily adjust the β-pairing in two well-aligned β-strands. The eighth auxiliary movement is β-turn formation, which attempts to form a β-turn in regions of the protein whose input secondary structure is defined as coiled. The final two movements are β-strand and α-helix formation. For these two movements, two regions that are defined as β-strands or α-helices are moved closer together based on distance and torsion angle distributions collected from the PDB.

**Text S2: FoldDesign energy function.**

The energy function used to guide the FoldDesign simulations is a combination of 10 energy terms:

$$E_{DeepFold} = E_{HB} + E_{ss\_satisfaction} + E_{rama} + E_{hhpack} + E_{sspack} + E_{\mathrm{hspack}} + E_{ev} + E_{generic\_dist} + E_{frag\_dist\_profile} + E_{frag\_solv} + E_{rg} + E_{contact\_num} \quad (1)$$

where $E_{HB}$, $E_{ss\_satisfaction}$, $E_{rama}$, $E_{hhpack}$, $E_{sspack}$, $E_{\mathrm{hspack}}$, $E_{ev}$, $E_{generic\_dist}$, $E_{frag\_dist\_profile}$, $E_{frag\_solv}$, $E_{rg}$, and $E_{contact\_num}$ are terms for backbone hydrogen bonding, secondary structure satisfaction, Ramachandran torsion angles, helix-helix packing, strand-strand packing, helix-strand packing, excluded volume, generic backbone atom distances, fragment-derived distance restraints, fragment-derived solvent accessibility, radius of gyration, and expected contact number, respectively. The equations for each energy term are detailed below.

$E_{HB}$ is calculated as follows:

$$E_{HB} = \sum_{i,j,T_k} E_{hb\_feat}(i,j,T_k)$$

where $i$ and $j$ are the residue indices and $T_k$ is the k$^{th}$ type of hydrogen bonding restraint. In FoldDesign, there are 4 types of hydrogen bonding restraints: hydrogen bonds between residues $i$ and $i+4$ in regions defined as helical by the input secondary structure ($T_1$), virtual hydrogen bonds between residues $i$ and $i+3$ in regions defined as helical by the input secondary structure ($T_2$), and hydrogen bonds between residues $i$ and $j$ in parallel β-strands ($T_3$) or antiparallel β-strands ($T_4$) for regions defined as strands by the input secondary structure. The energy for each type of hydrogen bonding restraint is calculated using the following equation:

$$E_{hb_{feat}}(i,j,T_k) = \sum\nolimits_{l=1}^{n_k} \frac{(f_l(i,j) - \mu_{kl})^2}{2\delta_{kl}^2}, \quad n_k = \begin{cases} 4 & k = 1,2 \\ 3 & k = 3,4 \end{cases}$$

where $f_l(i,j)$ is the value of the $l^{th}$ feature from the decoy structure, $n_k$ is the number of features considered for the $k^{th}$ type of hydrogen bond restraint, $\mu_{kl}$ is the average value of the $l^{th}$ feature for the $k^{th}$ type of hydrogen bond restraint calculated from the PDB library, and $\delta_{kl}$ is the standard deviation of the $l^{th}$ feature for the $k^{th}$ type of hydrogen bond restraint. For hydrogen bonding, we consider four features: the distance, $D(O_i,H_j)$, between backbone atom $O_i$ from residue $i$ and the backbone hydrogen, $H_j$, from residue $j$, the angle, $A(C_i,O_i,H_j)$, between backbone atoms $C_i$ and $O_i$ from residue $i$ and the backbone hydrogen, $H_j$, from residue $j$, the angle, $A(C_i,O_i,H_j)$, between backbone atom $O_i$ from residue $i$ and the backbone hydrogen, $H_j$, and nitrogen, $N_j$, from residue $j$, and the torsion angle, $T(C_i,O_i,Hj,N_j)$, between atoms $C_i$ and $O_i$ from residue $i$ and the backbone hydrogen, $H_j$, and nitrogen, $N_j$, from residue $j$. Note for hydrogen bonding in strand regions, $T_3$ and $T_4$ restraints, $T(C_i,O_i,Hj,N_j)$ is not considered as there is a large standard deviation for this feature in strand regions. The values of $\mu_{kl}$ and $\delta_{kl}$ are shown in Table S3.

$E_{ss\_satisfaction}$ is calculated as follows:

$$E_{ss\_satisfaction} = -\sum_{i=1}^{i=L}\begin{cases} -2 & if\ ss_i = helix\ and\ input\_ss_i = strand\ or\ ss_i = strand\ and\ input\_ss_i = helix \\ 1 & if\ ss_i = coil\ and\ input\_ss_i = coil \\ 2 & if\ ss_i = helix\ and\ input_{ss_i} = helix\ or\ ss_i = strand\ and\ input_{ss_i} = strand \\ -1 & else \end{cases}$$

where $ss_i$ is the secondary structure of the decoy at position $i$ and $input\_ss_i$ is the input secondary structure at the corresponding position. If the input secondary structure is defined as helical and the secondary structure of the decoy structure is a strand or if the input secondary structure is defined as a strand and the secondary structure of the decoy structure is helical, then a penalty of -2 is assigned to penalize opposite secondary structure assignments more heavily. Similarly, if the helical or strand regions are correct in the decoy structure, then a stronger bonus is assigned. Mismatches in coiled regions are penalized less heavily, and correctly generated coiled regions are also rewarded to a lesser degree as they are more flexible and lack regular hydrogen bonding patterns.

$E_{rama}$ is calculated as follows:

$$E_{rama} = -\sum_{i=2}^{i=L-1}\log\left(P(\phi_i,\psi_i) \mid input\_ss_i\right)$$

where $\phi_i$ and $\psi_i$ are the backbone torsion angles at position $i$ and $input\_ss_i$ is the input secondary structure at position $i$. The probabilities for each backbone torsion angle pair were determined from the I-TASSER (4) PDB library based on the secondary structure at a given position.

$E_{hhpack}$, $E_{sspack}$, and $E_{hspack}$ are calculated as follows:

$$E_{hhpack} = -\sum_{i,j}\log\left(P\left(\psi_{ij},\theta_{ij},\Phi_{ij}\right) \mid seq\_sep\right) - \sum_{i,j}\log\left(P\left(D_{ij},\theta_{ij}\right) \mid seq\_sep\right)$$

$$E_{sspack} = -\sum_{i,j}\log\left(P\left(\psi_{ij},\theta_{ij},\Phi_{ij}\right) \mid seq\_sep\right) - \sum_{i,j}\log\left(P\left(D_{ij},\theta_{ij}\right) \mid seq\_sep\right)$$

$$E_{hspack} = -\sum_{i,j} \log\big(P\big(\psi_{ij}, \theta_{ij}, \Phi_{ij}\big) \,\big|\, seq\_sep\big) - \sum_{i,j} \log\big(P\big(D_{ij}, \theta_{ij}\big) \,\big|\, seq\_sep\big)$$

where $\psi_{ij}$, $\theta_{ij}$, $\Phi_{ij}$ are the angles between two secondary structure elements (either two helices, $E_{hhpack}$, two strands $E_{sspack}$, or a helix and a strand, $E_{hspack}$) defined in Fig. S5, $D_{ij}$ is the distance between the centers of the two secondary structure elements, and $seq\_sep$ is the number of residues between two secondary structure elements. The potential is split into three different groups depending on the sequence separation, including short, medium, and long-range interactions. Here, short, medium, and long-range refers to residue pairs (*i,j*) that fall in the following ranges, respectively: $6 \le |i-j| < 12$, $12 \le |i-j| < 24$, and $|i-j| \ge 24$. The probabilities distributions for the features were derived from PDB structures in the I-TASSER library and were fit using kernel density estimation to smoothen the potentials.

$E_{ev}$ is calculated as follows:

$$E_{ev} = \sum_{i=1}^{i=L} \sum_{j=i+1}^{j=L} \sum_{ii} \sum_{jj} \begin{cases} (vdw(i,ii) + vdw(j,jj))^2 - r_{ii,jj}^2 & if\ r_{ii,jj} < vdw(i,ii) + vdw(j,jj) \\ 0 & else \end{cases}$$

where clashes are calculated between each atom ii from residue $i$ and atom $jj$ from residue $j$ and $r_{ii,jj}$ is the distance between the two atoms. For the side-chain center atoms, the center of mass of valine is used to assess steric clashes. All atoms presented in Fig. S2 are considered except for hydrogen.

$E_{generic\_dist}$ is calculated as follows:

$$E_{generic\_dist} = \sum_{i=1}^{i=L} \sum_{j=i+1}^{j=L} \sum_{ii} \sum_{jj} -RT * \log\left(\frac{N_{obs}(ii,jj,r_{ii,jj})}{r_{ii,jj}^{\alpha} N_{obs}(ii,jj,r_{cut})}\right)$$

where *L* is the protein length, *i* and *j* are the two residue indices and *ii/jj* are the atoms N, Cα, C, O and Cβ. $N_{obs}(ii,jj,r_{ii,jj})$ is the observed number of pairs between atoms *ii* and *jj* with distance $r_{ii,jj}$ determined from the I-TASSER PDB library. A cutoff, $r_{cut}$, of 15Å is used and the distances for the observed atom pairs is divided into 0.5Å bins from 0Å to 15Å. The potential is similar to DFIRE, where $\alpha = 1.61$ and $N_{obs}(ii,jj,r_{cut})$ is used to calculate the background probability.

$E_{frag\_dist\_profile}$ is calculated as follows:

$$E_{frag\_dist\_profile} = -\sum_{(i,j)\subseteq S_{dp}} \log\left(N_{ij}(d_{ij})\right)$$

where $d_{ij}$ is the distance between the Cα atoms of residues *i* and *j* in the decoy structure and $N_{ij}$ is the distance profile for residues *i* and *j* extracted from the 10 residue long fragments where *d* falls in the range [0Å, 9Å] with a bin width of 0.5 Å. $S_{dp}$ is the set of residues that have fragment-derived distance profiles. To derive the distance profiles, we first analyze each of the 10 residue fragments that originate from the same PDB structure and are aligned to different residues, *i* and *j*. Then we calculate the distance between the Cα atoms for the two positions from the fragments based on their corresponding positions in their PDB structure. If the distance between the two residues in the PDB structure is <9Å, then these positions may be encouraged to from contacts in the designed structure. This procedure is repeated for each query residue pair (*i*, *j*) to construct a histogram of distances. If the histogram for a given pair of residues has a peak <9Å, then the histogram is saved to calculate the distance profile energy and the residue pair is added to the set $S_{dp}$.

$E_{frag\_solv}$ is calculated as follows:

$$E_{frag\_solv} = \sum_{i=1}^{i=L} \left| s_i - s_i^E \right|$$

where $L$ is the protein length, $s_i$ is the solvent accessibility of residue *i* in the decoy structure, and $s_i^E$ is the expected solvent accessibility derived from the 20 residue fragments. The following formula is used to calculate $s_i$:

$$s_i = 1 - 0.007 \sum_{d(G_i,G_j)<9\text{Å}} \frac{A_{aa(j)}}{d^2(G_i, G_j)}$$

Here, $A_{aa(j)}$ is the maximum solvent accessible surface area for the given residue $aa$ at position *j*. Since polyvaline sequences are used in FoldDesign, the maximum solvent accessible surface area for Valine is used. $G_i$ and $G_j$ are the geometric centers of residues *i* and *j*, $d(G_i, G_j)$ is the distance between the two geometric centers, and $d^2(G_i, G_j)$ is the squared distance. A cutoff of 9Å is used as residues that are further apart contribute little to the solvent accessibility. As mentioned above, $s_i^E$ is the expected solvent accessibility calculated from the overlapping 20 residue fragments. For each fragment, the solvent accessibility of the residue in its native PDB structure is recorded, and the estimated solvent accessibility is calculated by averaging the solvent accessibility of each fragment residue aligned to position *i*.

$E_{rg}$ is calculated as follows:

$$E_{rg} = \begin{cases} 0 & r_{min} \le r \le r_{max} \\ (r_{min} - r)^2 & r < r_{min} \\ (r - r_{max})^2 & r > r_{max} \end{cases}$$

where $r$ is the radius of gyration for the decoy structure calculated from the Cα positions produced during the FoldDesign simulations and $r_{min}$/$r_{max}$ are the estimated minimum and maximum radii of gyration calculated from the PDB based on the protein length and secondary structure composition.

$E_{contact\_num}$ is calculated as follows:

$$E_{contact\_num} = |num_{short\_cont} - expected\_num_{short\_cont}| + |num_{med\_cont} - expected\_num_{med\_cont}| + |num_{long\_cont} - expected\_num_{long\_cont}|$$

where $num_{short\_cont}$, $num_{med\_cont}$, and $num_{long\_cont}$ are the number of short, medium, and long-range contacts in the decoy structure. Here, short, medium, and long-range contacts refer to residue pairs (*i,j*) that fall in the following ranges, respectively: $6 \leq |i-j| < 12$, $12 \leq |i-j| < 24$, and $|i-j| \geq 24$. $expected\_num_{short\_cont}$, $expected\_num_{med\_cont}$, and $expected\_num_{long\_cont}$ are the expected short, medium, and long-range contacts calculated from PDB structures in the I-TASSER library based on protein length.

**Text S3: Rosetta protocol used to generate designed folds.**

The following command was used to generate backbones by Rosetta:

```
<rosetta_bin>/main/source/bin/rosetta_scripts.static.linuxgccrelease -database <rosetta_bin>/main/database/ -s ./input.pdb -parser:protocol ./backbone_generation.xml -nstruct 250 -overwrite
```

For each input topology, 250 designs were generated using Rosetta, where the final designs were selected from the lowest energy designs as assessed by the Rosetta centroid energy function. The contents of the backbone_generation.xml files are detailed below, which were adapted from a representative recent publication (5).

```
<ROSETTASCRIPTS>
    <SCOREFXNS>
      <ScoreFunction name="SFXN1" weights="fldsgn_cen_omega02.wts" />
    </SCOREFXNS>
    <FILTERS>
        <ScoreType name="cen_total" scorefxn="SFXN1" score_type="total_score" threshold="1000000" />
        <ScoreType name="vdw" scorefxn="SFXN1" score_type="vdw" threshold="1000000" />
        <ScoreType name="rg" scorefxn="SFXN1" score_type="rg" threshold="1000000" />
        <ScoreType name="cen_rama" scorefxn="SFXN1" score_type="rama" threshold="1000000" />
        <ScoreType name="sspair" scorefxn="SFXN1" score_type="ss_pair" threshold="1000000" />
        <ScoreType name="rsigma" scorefxn="SFXN1" score_type="rsigma" threshold="1000000" />
    </FILTERS>
    <TASKOPERATIONS>
    </TASKOPERATIONS>
    <MOVERS>
<Dssp name="dssp"/>
        <SwitchResidueTypeSetMover name="fullatom" set="fa_standard"/>
        <SwitchResidueTypeSetMover name="cent" set="centroid"/>
        <MakePolyX name="polyval" aa="VAl" keep_pro="1" />
        <BluePrintBDR name="bdr1" scorefxn="SFXN1" use_abego_bias="1" blueprint="blueprint.xml"/>
        <MinMover name="min1" scorefxn="SFXN1" chi="1" bb="1" type="dfpmin_armijo_nonmonotone_atol"
tolerance="0.0001"/>
        <MinMover name="cart_min1" scorefxn="SFXN1" type="lbfgs_armijo_nonmonotone" tolerance="0.0001"
max_iter="1000" chi="0" bb="1" bondangle="1" bondlength="1" cartesian="1"/>
        <ParsedProtocol name="cenmin1" >
          <Add mover_name="cent" />
          <Add mover_name="min1" />
          <Add mover_name="fullatom" />
        </ParsedProtocol>
        <ParsedProtocol name="bdr1ss" >
          <Add mover_name="bdr1" />
          <Add mover_name="cenmin1" />
```

```
        <Add mover_name="dssp" />
      </ParsedProtocol>
  </MOVERS>
  <PROTOCOLS>
      <Add mover_name="bdr1ss" />
      <Add mover_name="fullatom" />
      <Add filter_name="cen_total" />
      <Add filter_name="vdw" />
      <Add filter_name="rg" />
      <Add filter_name="cen_rama" />
      <Add filter_name="sspair" />
      <Add filter_name="rsigma" />
  </PROTOCOLS>
</ROSETTASCRIPTS>
```

The contents of the weights file (fldsgn_cen_omega02.wts) were as follows, which were also adapted from the previous study (5):

```
vdw 1.0
rg 1.0
rama 0.1
hs_pair 1.0
ss_pair 1.0
rsigma 1.0
omega 0.5
hbond_lr_bb 1.0
hbond_sr_bb 1.0

STRAND_STRAND_WEIGHTS 1 11
```
